# One-year descriptive analysis of patients treated at an anti-rabies clinic – a retrospective study from Kashmir

**DOI:** 10.1101/644393

**Authors:** Khalid Bashir, Inaamul Haq, S Muhammad Salim Khan, Mariya Amin Qurieshi

## Abstract

**Background:** Dog bites in humans are a major public health problem in India in general and Kashmir in particular. Canine rabies is almost non-existent in developed countries and exists mainly in the poorer, low socioeconomic strata of society in the developing world. The objective of this study was to determine the characteristics, pattern, and burden of dog bite injuries in the Kashmir valley.

**Methodology and principal findings:** Data from Anti-Rabies Clinic of a tertiary care hospital in Srinagar, the summer capital of the state of Jammu & Kashmir, was collated and analyzed. Analysis of records of all the patients who had reported between April 2016 and March 2017 was done. A total of 6172 patients had reported to the Anti-Rabies Clinic for management of animal bites from 1^st^ April 2016 to 31^st^ March 2017. Most of the patients were young males. Almost half (47.7%) of the patients were bitten in the afternoon. Lower limbs were the most common site of bite (71.7%). Most of the bites were of Category III (57.6%) followed by Category II (42.3%); only one case of Category I was recorded. Almost all (98.0%) cases reported being bitten by dogs.

**Conclusions:** Category III dog bites on lower limbs are the most common type of animal bites. Children have more chances of a bite on head and neck region. Serious and workable efforts have to be made to reduce the incidence and consequences of animal bites.

**Author summary:** In Kashmir, dog bite is an important public health problem. Thousands of people become victims of an animal bite, especially dog bite, and some of them develop rabies. Rabies is an invariably fatal viral disease resulting in approximately 59 000 human deaths per year globally, with 95% of cases occurring in Africa and Asia. The only way to prevent a rabies death is vaccination of an animal bite victim. In Kashmir, the burden and characteristics of dog bites are not routinely captured by the health system in place. We, therefore, attempted to find out the burden and characteristics of animal bite victims by analyzing one-year data from an Anti-Rabies Clinic at a tertiary care hospital in Kashmir. We found that 98% of the patients registered at the clinic during the period were victims of a dog bite. The victims were mostly young males but females and children were not shown any mercy either. Lower limbs were the most favorite site of the bite. Our analysis of the data also revealed that children under 15 years were more prone to a bite in the head and neck region. We concluded that the burden of animal bites, especially dog bites, is huge in Kashmir and recommended that serious efforts directed towards immunizing and decreasing the stray dog population need to be put into practice to decrease the number of animal bite victims and prevent any rabies deaths.

## Introduction

Rabies transmission by dog bites is one of the major public health problems, considering its palpable fear and anxiety, as it is a sinister zoonotic infection transmitted to humans or animals by the bite of a rabid animal. Rabies virus, which belongs to the Lyssavirus genus of the *Rhabdoviridae* family, causes fatal encephalitis [1]. It imposes high economic costs annually in various countries due to its high incidence [2,3]. Dog bites constitute more than 95% of human rabies cases and the rest is associated with a cat, fox and other carnivores in developing countries. A bite by a potentially rabid dog represents the source of infection in more than 99% of cases [4,5]. Among all PEP recipients, 30-50% are children <15 years of age and they also have the highest rabies mortality [6,7]. Dog rabies is almost non-existent in developed countries of Europe, North America, and Australia; but it is prevalent in the developing world. Lack of reliable data and unawareness of the burden and risk factors associated with human rabies together represent a critical challenge for the formulation of policies and strategies to control the disease and has been considered a major cause for underinvestment in rabies control measures in these countries [8]. In developing countries, the real number of sufferers is probably higher than the reported statistics [9]. In addition to the health importance in human-beings, disease outbreak among live-stock causes significant economic losses. Despite the preventability of rabies by effective and safe vaccines, the disease is still a healthcare problem in many countries, especially in Africa, Asia, and India [10]. Asia constitutes 96.5% of the burden of the disease in developing countries, which costs it 560 million USD annually [11,12].

Rabies is invariably fatal, yet completely preventable if post-exposure prophylaxis (PEP) is applied in a timely and correct manner. The World Health Organization (WHO) has prepared standard recommendations for PEP, which include immediate and thorough wound washing with soap and water, followed by administration of the vaccine and additionally, infiltration of Rabies Immunoglobulin in WHO category III bites [13].

In Kashmir, there is a huge and ever-increasing population of free-roaming stray dogs. The present study aims to generate a picture of the alarmingly increasing dog bite victims, potential cases of dreaded rabies, whose quality of physical and psychological health is directly or indirectly influenced by the event. Our objective was to determine the epidemiological features, characteristics, and pattern of dog bite victims who received PEP at a tertiary care center in Kashmir for one year. This will help in devising prevention strategies in view of the elimination of dog-mediated human rabies by 2030, as jointly outlined by WHO, FAO, OIE, the Global Alliance for Rabies Control, and the international community [14].

## Methods

This study was conducted at the Anti-Rabies Clinic (ARC) of Shri Maharaja Hari Singh (SMHS) Hospital, Government Medical College, Srinagar, run by the Department of Community Medicine. The ARC receives animal bite cases from the whole of Kashmir valley which had a population of 6.9 million as per the 2011 census [15]. The ARC maintains records of demographic and clinical details of patients visiting the clinic for treatment. The clinic adheres to WHO-recommended protocol for PEP, which includes prompt wound washing, an anti-rabies vaccine for WHO Category II and III exposures, and use of Immunoglobulin in Category III exposures [13]. Following recommendations of the World Health Organization on PEP following animal bites in 2005, the ARC started the use of the intradermal regimen in 2011. Recently the ARC has started the 2-site intradermal schedule as recommended by the recent WHO document [16].

The ARC follows a protocol for animal bite management which includes tetanus toxoid immunization for category II and III bites and Equine Rabies Immunoglobulin (ERIG) 40 IU/Kg body weight for category III bites. The ARC provides anti-rabies vaccines free of cost to the bite victims. ERIG is not provided free of cost. Patients need to purchase ERIG from the market. The health workers at the ARC are trained in the management of animal bites as well as the administration of the vaccine and ERIG.

We did an analysis of secondary data; records of the patients from 1^st^ April 2016 to 31^st^ March 2017 were collated and analyzed. All data analyzed were anonymized. Demographic and clinical details of the patients were entered into a Microsoft Excel spreadsheet. From records, we took into consideration different variables for study analysis including age, sex, time, site, severity and WHO category of bite, type of animal involved and the residence of the patient.

### Statistical analysis

Data was entered into a Microsoft Excel spreadsheet. Chi-square test was used to test independence between the site of bite versus age and sex. In order to further explore the relationship between site of bite and sex, analysis restricted to cases >15 years of age was done. Chi-squared goodness-of-fit was used to test if the number of cases varied across months. A p-value of less than 0.05 was considered statistically significant. SPSS version 23.0 (IBM Corp. Released 2015.IBM SPSS Statistics for Windows, Version 23.0. Armonk, NY: IBM Corp.) was used for analyzing the data.

## Results

A total of 6172 patients had reported to the ARC from 1^st^ April 2016 to 31^st^ March 2017. The demographic details of the patients are shown in Table 1. The bite victims were mostly young and middle-aged males.

**Table 1:** Demographic characteristics of animal bite victims

| Demographic characteristics | Number of bite victims (n=6172) | Percentage (%) |
| --- | --- | --- |
| **Age (years)** |  |  |
| ≤5 | 426 | 6.9 |
| 6-15 | 986 | 16.0 |
| 16-30 | 1519 | 24.6 |
| 31-50 | 2278 | 36.9 |
| 51-65 | 773 | 12.5 |
| >65 | 190 | 3.1 |
| **Sex** |  |  |
| Female | 1635 | 26.5 |
| Male | 4537 | 73.5 |
| **Residence** |  |  |
| District Srinagar | 3773 | 61.1 |
| Other districts | 2399 | 38.9 |

The number of cases was highest during the months of March to May (Fig 1).

**Fig 1.**
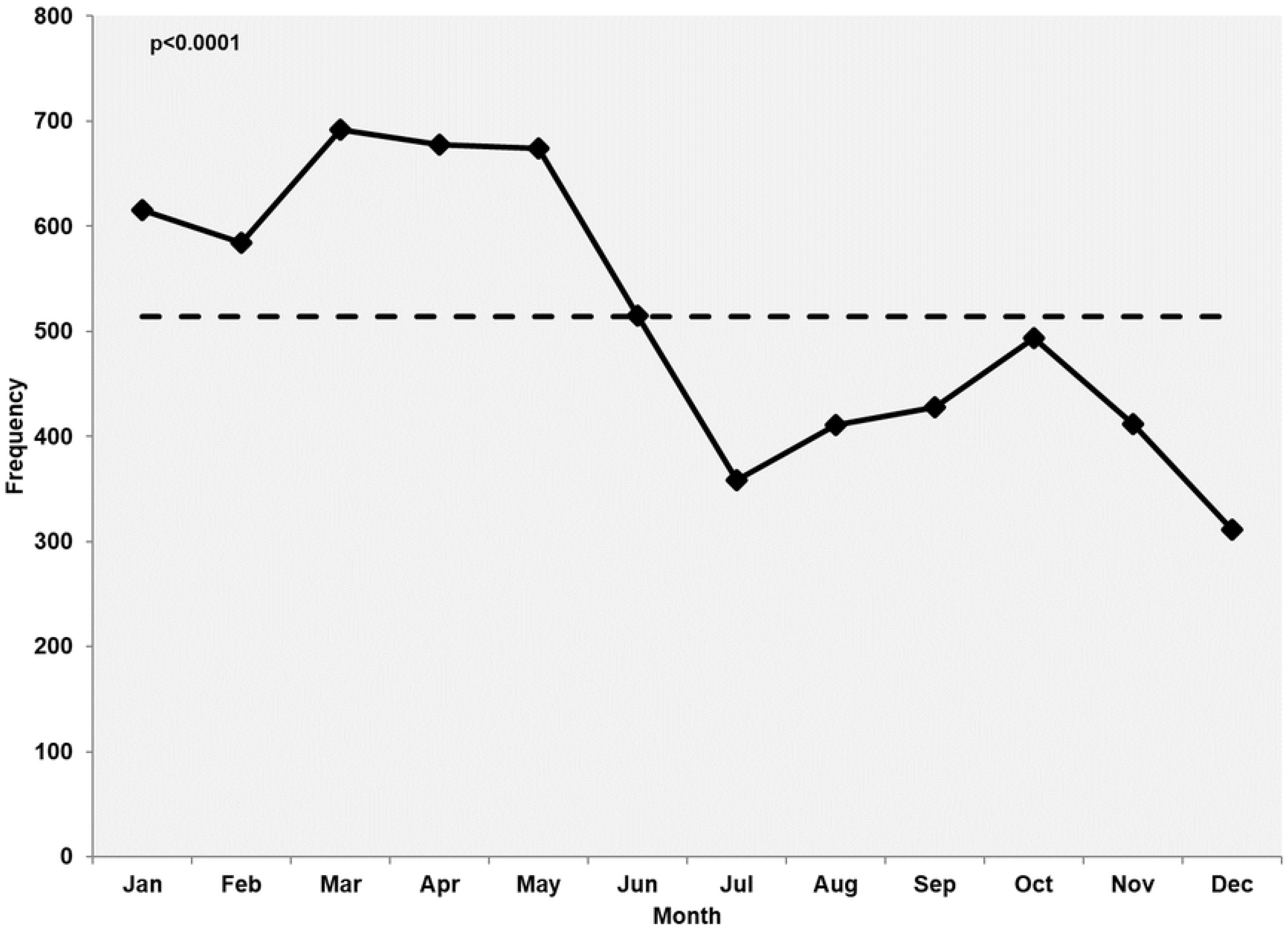
Monthly distribution of animal bite cases. Dash line indicates the monthly average of animal bite cases. The cases were the highest during the months following the winter months.

Table 2 describes the characteristics of animal bites. Almost half of the cases (47.7%) were bitten in the afternoon (1200 hours to 1759 hours). Lower limbs were the most common site of bite (71.7%). Ninety-eight percent of patients were bitten by a dog (6048/6172). Only one patient reported with a WHO category I bite.

**Table 2:**
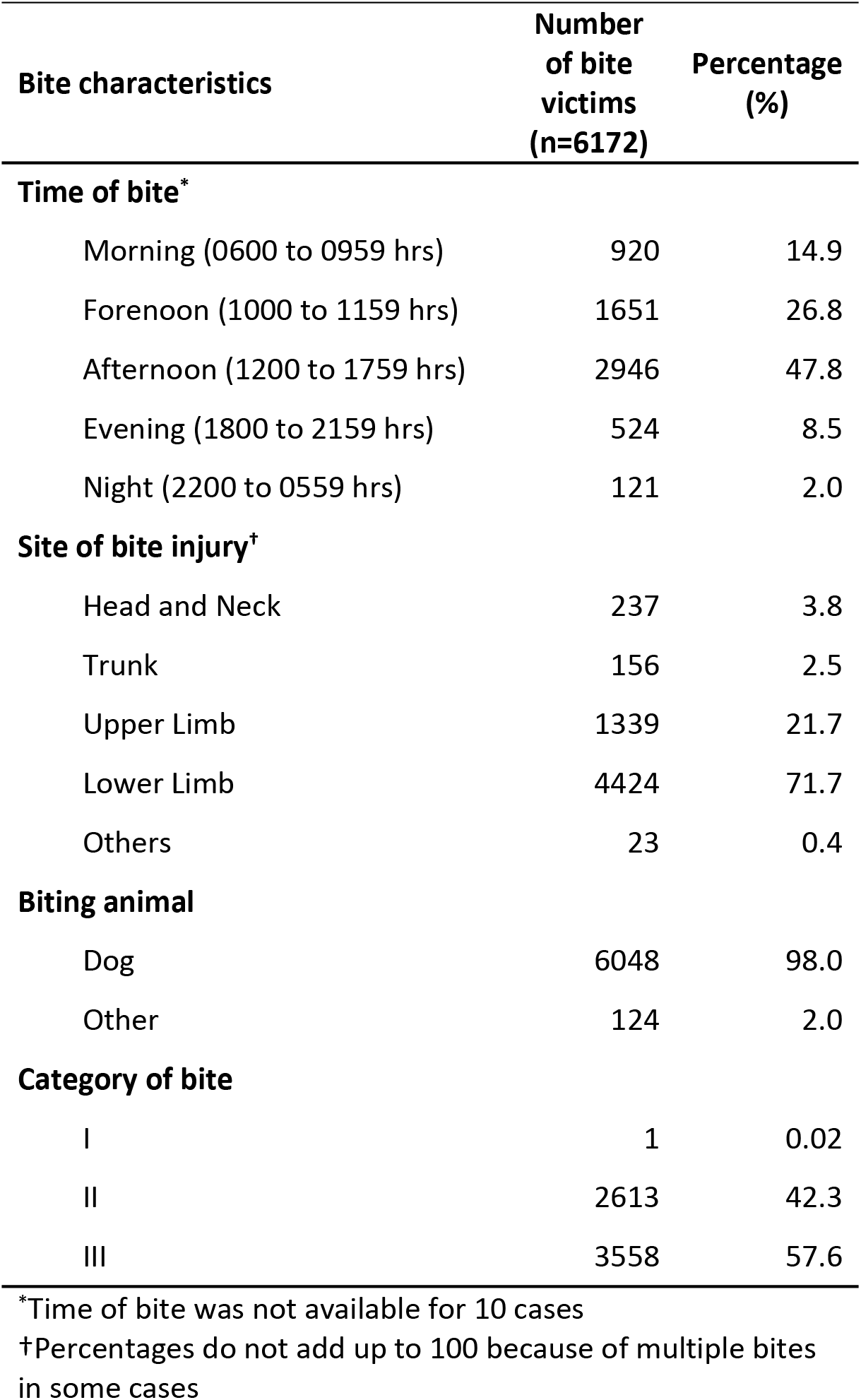
Characteristics of animal bites

| Bite characteristics | Number of bite victims (n=6172) | Percentage (%) |
| --- | --- | --- |
| **Time of bite\*** |  |  |
| Morning (0600 to 0959 hrs) | 920 | 14.9 |
| Forenoon (1000 to 1159 hrs) | 1651 | 26.8 |
| Afternoon (1200 to 1759 hrs) | 2946 | 47.8 |
| Evening (1800 to 2159 hrs) | 524 | 8.5 |
| Night (2200 to 0559 hrs) | 121 | 2.0 |
| **Site of bite injury†** |  |  |
| Head and Neck | 237 | 3.8 |
| Trunk | 156 | 2.5 |
| Upper Limb | 1339 | 21.7 |
| Lower Limb | 4424 | 71.7 |
| Others | 23 | 0.4 |
| **Biting animal** |  |  |
| Dog | 6048 | 98.0 |
| Other | 124 | 2.0 |
| **Category of bite** |  |  |
| I | 1 | 0.02 |
| II | 2613 | 42.3 |
| III | 3558 | 57.6 |

\*Time of bite was not available for 10 cases
†Percentages do not add up to 100 because of multiple bites in some cases

Table 3 analyzes the relationship between the site of bite versus age and sex. The site of the bite was significantly associated with age (p<0.0001). Lower limbs were the most common site of bite across all age and sex groups. However, under-five children were more likely to be bitten on the head, neck and trunk - adjusted standardized residual (ASR) 8.3 and 8.2, respectively. Young adults were more likely to be bitten on lower limbs than expected (ASR 3.7). Males were more likely to be bitten on lower limbs (ASR 2.4) and females were more likely to be bitten on upper limbs (ASR 2.7) than expected.

**Table 3:**
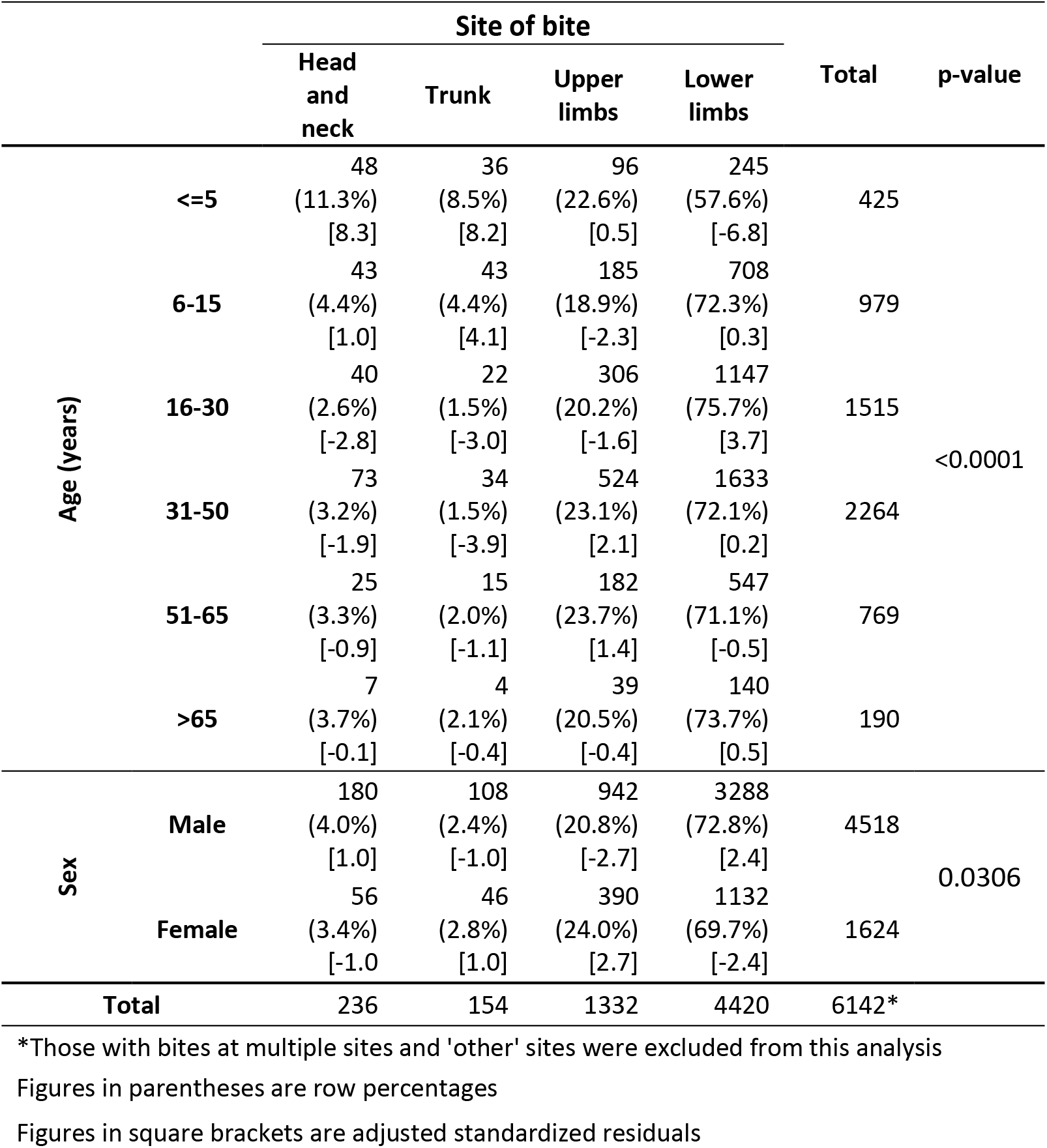
Relationship of site of bite versus age and sex

Analysis restricted to patients >15 years of age (Table 4) further revealed that as compared to females, males were more likely than expected to be bitten on the head and neck region (ASR 2.9).

**Table 4:**
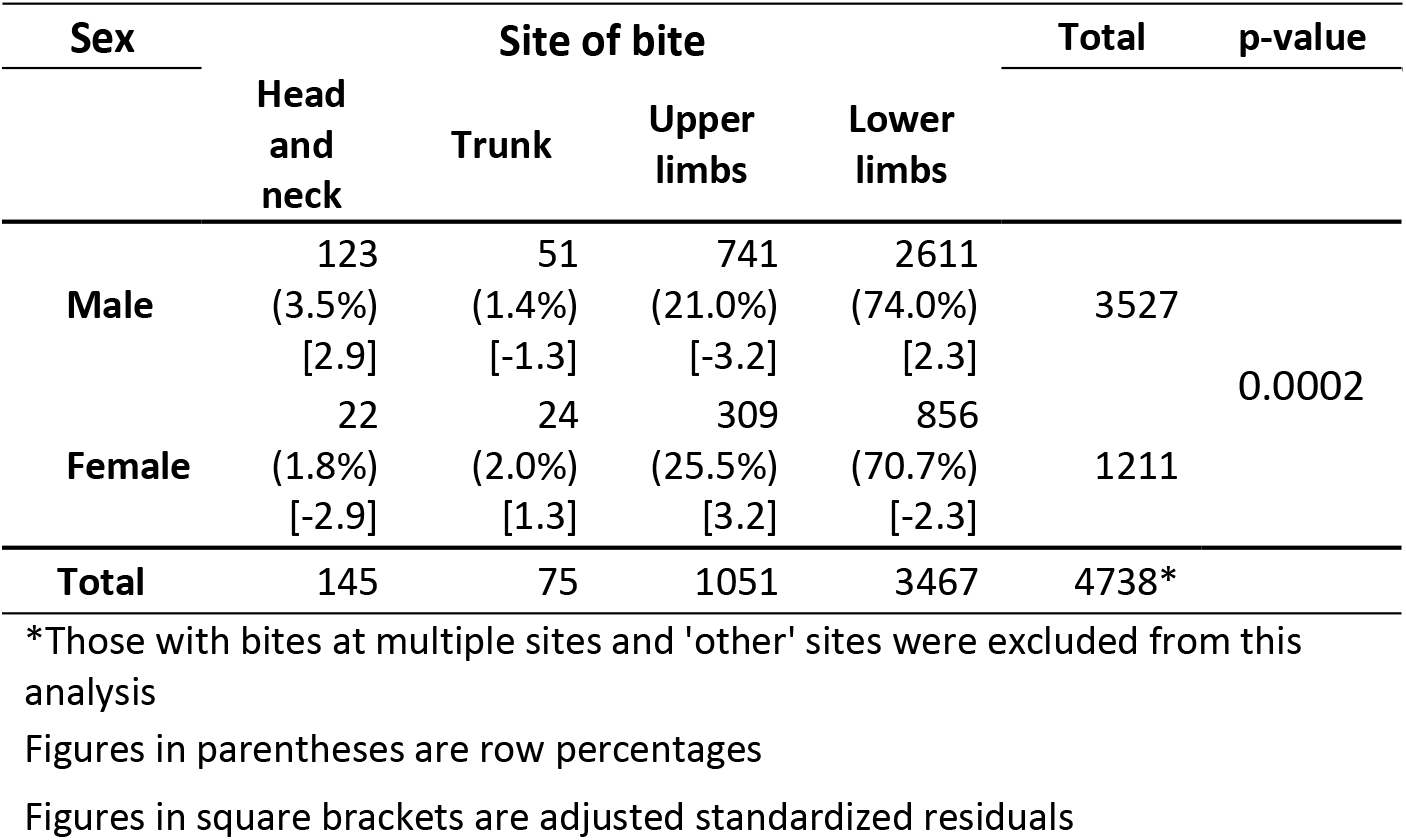
Sex versus site of bite in animal bite victims >15 years of age

## Discussion

This study shows that dog bite-related injuries are very common in Kashmir. Category III dog bites on the lower limbs are the most common.

We analyzed data from the only ARC in Kashmir which mostly provides services to people from District Srinagar. In our study, 3773 out of a total of 6172 cases were from Srinagar. Animal bite cases are being managed at other hospitals in the District as well. During the study period, 2032 animal bite cases were managed at these hospitals. This gives us a total of 5805 animal bite cases during the study period. We estimated the mid-interval population of Srinagar during the period of study to be 1290644 based on 2011 census data [16] for the district and the growth rates for urban areas of the state of Jammu & Kashmir reported by the Sample Registration System [17]. Thus, the estimated incidence of an animal bite in Srinagar during the one-year study period was 450 per 100,000 population. This is much higher than the incidence reported from Kenya [18] (284 per 100,000), Tanzania [19] (74 per 100,000) and Ghana [20] (54.1 per 100,000).

The cases were mostly middle-aged men (Table 1). But children and elderly were not spared. The youngest victim was an infant and the oldest victim was 97 years old. About 23% were children <15 years of age. Males outnumbered females. In Kashmir, males usually go out more frequently for work or for social visits as compared to females. Most of the available literature on dog bites has reported a male predominance [18,19,21–24]. The percentage of child victims varies across studies from 25% to 72% [18–26].

The ARC is situated in District Srinagar and that might be the reason why most of the cases in our study were from Srinagar. District hospitals and some of the sub-district level hospital in other districts of the region provide bite-management services to people in those areas.

Our analysis shows a definite rise in the number of cases during the months of March to May (Fig 1). Kashmir is a valley with the winter season starting in December and ending in February. People usually prefer to stay indoors during the period. Outdoor movement of people increases as the winter ends. This might have led to increased interaction with dogs and hence an increase in dog bite cases during the months following winter. A somewhat similar trend has been reported from Iran [21]. However, in a five-year study from Srilanka, Kularatne et al reported a more or less even pattern across seasons [24].

The afternoon was the most common time of animal bites (47.7%) followed by forenoon (26.7%) (Table 2). Every seventh case was bitten when he/she ventured out early morning, which people usually do for the morning prayers or for a visit to the bakery or some other work. The most common site of the bite was lower limbs followed by upper limbs (Table 2). Seven patients had bites at more than one region. Consumption of milk from a rabid cow was the most common among the “others” category. Our results are similar to results from Kenya [18] and Nigeria [23]. Interestingly, an earlier study from Jos Plateau State of Nigeria [22] reported that 85% of victims were bitten on arms.

We found a significant relationship between a victim’s age and the site of an animal bite (p<0.0001) (Table 3). Lower limbs were the most common site of bite across all age and sex groups. A breakdown of table 3 using adjusted standardized residuals revealed that under-five and young children were more likely to be victims of an animal bite on the head, neck, and trunk.

Our results revealed a significant relationship between sex and the site of an animal bite (p=0.0308, Table 3). In order to further delve into the relationship, we restricted our analysis to patients >15 years of age (Table 4) hypothesizing that any sex differentials in this context will come into play after puberty. To confirm our notion, we stratified the analysis by age. Among children up to 15 years of age, there was no significant relationship between sex and the site of the animal bite. The analysis revealed an interesting relationship. As compared to females, males were more likely to report with a bite in the head and neck region. Furthermore, upper limbs were a more likely site for an animal bite among females.

A dog was the biting animal in 98% of cases. Records lacked information about dog ownership. However, our experience with dog bites at the ARC has been that most of the cases are due to street dogs as people in the region do not usually keep a dog as a pet. Other animals included leopard, cat, and cow.

More than half of the cases (57.6%) were classified as category III bites and only one category I bite was recorded.

Animal bite victims suffer huge losses in the form of lost wages, travel costs, and direct treatment costs. Moreover, psychological and emotional denting is something which is difficult to translate. Scarring is a common consequence related to dog bites, and the resulting emotional distress due to cosmetic reasons should not be undermined, particularly for wounds on the face.

The biggest challenges in Africa and Asia, including Kashmir in particular, are free-roaming dog populations, limited veterinary and human health infrastructure, and absence of efficient communication between the veterinary and the human health sectors [27,28]. The absence of effective control over the growing dog population is proving costly in terms of DALYs, premature death and the cost of PEP to public and private health sectors [28,29]. There are conflicting reports about the number of stray dogs in Srinagar with numbers varying from as low as 22,000 to as high as 1,50,000 [30–32]. Whatever the actual number of stray dogs, there is a dire need to devise a strategy to control the alarmingly growing dog population. Unfortunately, there is no provision in our region for isolating suspected dogs or for diagnosing animal rabies. Measures like mass dog vaccination campaigns to improve herd immunity, as recommended by veterinarians, can reduce the burden of human rabies [30]. Animal rabies control requires more attention and must be done through local municipalities. Rabies elimination programs focused mainly on mass vaccination of dogs are largely justified by the future savings of human rabies prevention programs.

### Limitations

This study was based on an analysis of records from an anti-rabies clinic. We, therefore, could not obtain information about the characteristics of the biting animal such as ownership or the circumstances of bite such as provocation. Information was not available about the receipt of tetanus vaccination and ERIG. Because of the secondary nature of the data, we could not validate the categorization of bite. However, during the study period bite categorization was done by trained health workers. The percentage of patients who completed PEP was also not available. We also could not evaluate the impact of animal bites on the physical and psychological health of the victims. An economic burden evaluation was also not possible. We could not draw the real picture of dog bite victims in Kashmir, as data analysis was limited to our hospital and many patients visit hospitals in the peripheries as well while some others miss PEP due to lack of knowledge.

### Conclusions

This study highlights the magnitude of animal bite injuries in Kashmir. Ninety-eight percent of patients had suffered injury from free roaming stray dogs. Category III bites on the lower limbs are the most common type of bite. Children are more prone to be bitten on the head, neck, and trunk. The burgeoning dog population takes a toll on quality of life in an explicit and implicit manner. It shakes up a person psychologically and emotionally which is often not perceived. Also, dog bites don’t spare children and geriatric population; in fact, they are easy prey. It is time to take proactive measures to stop the menace of the growing dog population.

## Supporting Information Legends

S1 Checklist: STROBE checklist

S2 Data: Anonymized data sheet

